# The Ca^2+^-activated Cl^-^ channel TMEM16B shapes the response time course of olfactory sensory neurons

**DOI:** 10.1101/2024.05.18.594801

**Authors:** Johannes Reisert, Simone Pifferi, Giorgia Guarneri, Chiara Ricci, Anna Menini, Michele Dibattista

## Abstract

Mammalian olfactory sensory neurons (OSNs) generate an odorant-induced response by sequentially activating two ion channels, which are in their ciliary membranes. First, a cationic, Ca^2+^-permeable cyclic nucleotide-gated channel is opened following odorant stimulation via a G protein-coupled transduction cascade and an ensuing raise in cAMP. Second, the increase in ciliary Ca^2+^ opens the excitatory Ca^2+^-activated Cl^-^ channel TMEM16B that carries most of the odorant-induced receptor current. While the role of TMEM16B in amplifying the response has been well established, it is less understood how this secondary ion channel contributes to response kinetics and action potential generation during single as well as repeated stimulation and, on the other hand, which response properties the CNG channel determines. We first demonstrate that basic membrane properties such as input resistance, resting potential and voltage-gated currents remained unchanged in OSNs that lack TMEM16B. The CNG channel predominantly determines the response delay and adaptation during odorant exposure, while the absence of the Cl^-^ channels shortens both the time the response requires to reach its maximum as well as to terminate after odorant stimulation. This faster response termination in *Tmem16b* knockout OSNs allows them, somewhat counterintuitively, to fire action potentials more reliably when stimulated repeatedly in rapid succession, a phenomenon that occurs both in isolated OSNs as well as in OSNs within epithelial slices. Thus, while the two olfactory ion channels act in concert to generate the overall response, each one controls specific aspects of the odorant-induced response.

## Introduction

Ethologically relevant odor-guided tasks, such as finding food, are crucial for animals to survive, failing to do so would cause death. Locating food based on its odor cues is dependent on a series of mechanisms starting at the very periphery of the olfactory system: odorant transduction mechanisms in olfactory sensory neurons (OSNs), which are located in the olfactory epithelium in the nasal cavity (Kleene, 2008; Tirindelli et al., 2009; Pifferi et al., 2012; Genovese et al., 2021). Inhaled odorants are detected by OSNs and are transduced first into a receptor current that then triggers action potentials that are conveyed to the olfactory bulb and secondary neurons (Cang and Isaacson, 2003; Spors et al., 2006; Gire et al., 2012; Tan et al., 2015). Olfactory transduction occurs in OSN cilia, which reach from the OSN dendrite into the mucus that lines the nasal cavity. Cilia contain the molecular machinery required for odor detection, which begins with the activation of an odorant receptor following odorant binding (Buck and Axel, 1991; Malnic et al., 1999). Adenylyl cyclase III (ACIII) (Bakalyar and Reed, 1990), via the G protein G_olf_ (Jones and Reed, 1989), increases cAMP which opens the olfactory cyclic nucleotide-gated (CNG) channel (Nakamura and Gold, 1987). This channel conducts Ca^2+^ and thus raises ciliary Ca^2+^ levels. Interestingly, Ca^2+^ then activates a Ca^2+^-activated Cl^-^ channel TMEM16B (also known as Anoctamin 2) (Kleene and Gesteland, 1991; Kurahashi and Yau, 1993; Lowe and Gold, 1993; Stephan et al., 2009; Hengl et al., 2010; Rasche et al., 2010; Sagheddu et al., 2010; Billig et al., 2011; Li et al., 2018). As ciliary Cl^-^ is high, this channel is excitatory and carries about 80 – 90 % of the transduction current, thus largely amplifying the initial CNG current (Kleene, 1993; Reisert et al., 2005; Boccaccio and Menini, 2007). Action potentials driven by the depolarizing transduction currents are typically only generated at the onset of the odorant-induced response with the amplitude of the action potentials quickly decreasing within an action potential train (Reisert and Matthews, 2001). This is most likely due to inactivation of voltage-gated Na^+^ and Ca^2+^ during the strong depolarization in OSNs, which have a very high input resistance (Trotier, 1994; Kawai et al., 1997). OSNs often only generate 2 – 3 action potentials when exposed to odorants (Reisert and Matthews, 2001).

The contribution of the amplitude and kinetics of odorant transduction to olfactory-driven behavior is not very well understood. Mouse models lacking G_olf_, ACIII, and the main subunit of the CNG channel, CNGA2, showed hardly any odorant responses, and pups had poor survival rates because of their inability to locate food and therefore feed (Brunet et al., 1996; Belluscio et al., 1998; Wong et al., 2000). The lack of response precluded more detailed analysis of their role in response kinetics. The last component in the series of events that lead to the generation of the transduction currents is the Ca^2+^-activated Clˉ channel TMEM16B and although its role in amplifying the receptor current is well established, its contribution to olfactory-driven behavior has been less clear cut. Originally it was concluded that TMEM16B might serve little to no role in olfactory behavior (Billig et al., 2011), with a later more nuanced evaluations of its role suggesting that it alters OSN action potential generation, axonal targeting of OSN axons to the olfactory bulb and bulbar activity, and odor-guided search tasks in particular for novel odors (Pietra et al., 2016; Neureither et al., 2017; Zak et al., 2018). Therefore, the role of the Cl^-^ channel in odor response magnitude and kinetics to shape OSN physiology, in particular in longer and repeated odor stimulation, and ultimately behavior is important to consider (Guarneri et al., 2023). In general, the kinetics of the peripheral odorant responses must be finely tuned to repeated odorant stimulations at breathing rates at rest or during high frequency sniff bouts (Ghatpande and Reisert, 2011). Therefore, understanding each aspect of the odorant response (i.e. termination, adaptation, action potential frequency and potential number as well as train duration) would help us to define the contribution of peripheral odor coding to overall olfactory coding. To tackle those points, we used a mouse model lacking TMEM16B to better understand response kinetics and odorant-induced adaptation. In addition, we designed OSN stimulation paradigms to mimic sniffing pattern to study how isolated OSNs generate action potentials as well as OSNs in slices of the olfactory epithelium.

## RESULTS

### The absence of Clˉ channels does not alter the basic biophysical properties of OSNs

We sought to understand whether the removal of the Clˉ channel would affect biophysical properties of OSNs. First, using the whole-cell patch clamp configuration, we recorded both inward and outward voltage-gated currents from OSNs in OE slices and found that they were similar in their amplitude and their voltage dependency between the *Tmem16b* KO and the WT (Fig. 1A-B). Then, we measured the input resistance and resting membrane potential of OSNs from WT and *Tmem16b* KO mice and found that those were not different (Fig. 1C-D).

**Figure 1.**
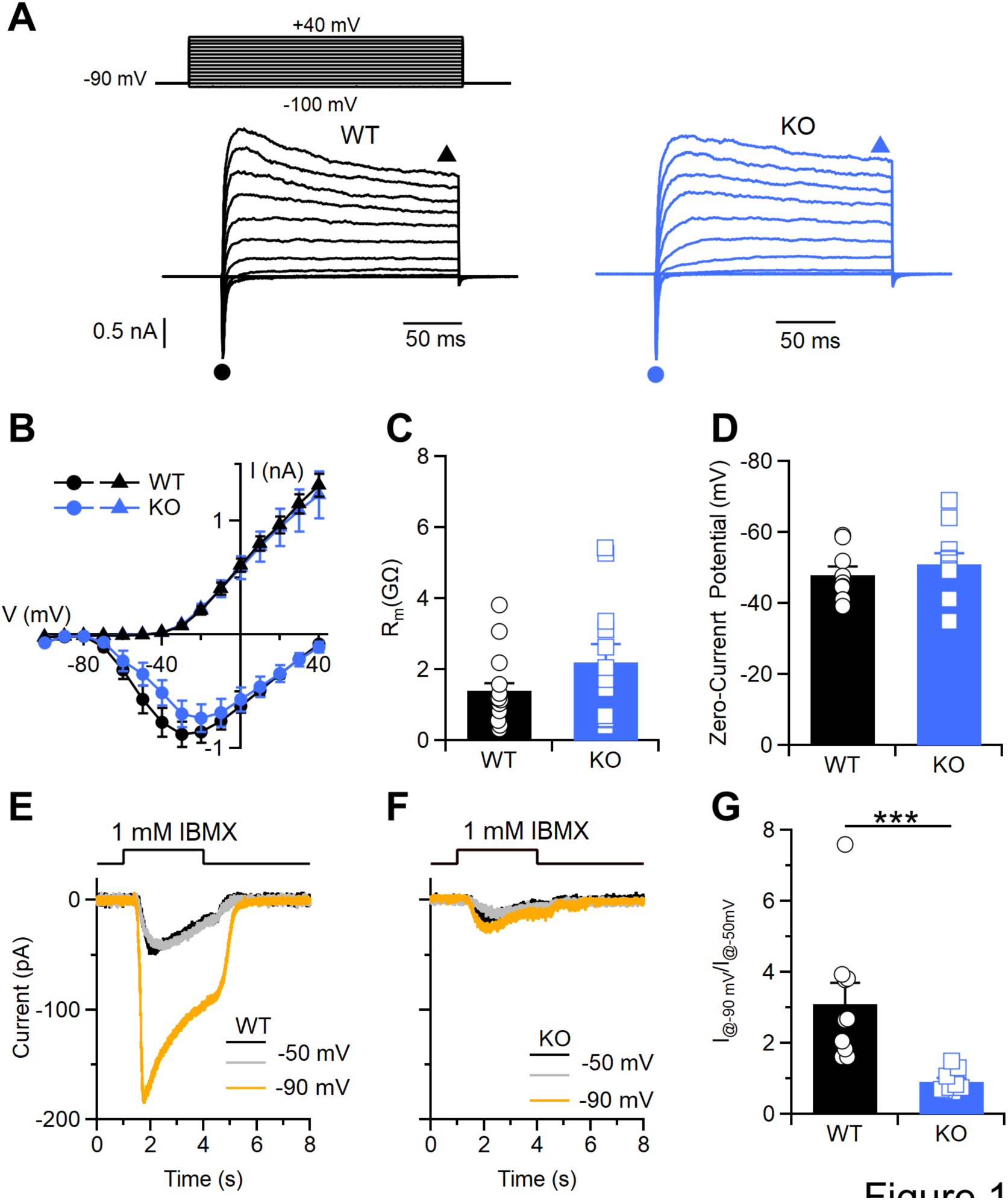
*Tmem16b* deletion abolishes the transduction Cl^-^ current without affecting the voltage-gated currents and membrane properties of OSNs. **A** Representative whole-cell recordings of OSNs from WT and *Tmem16b* KO mice. Voltage steps of 200-ms duration were given from a holding voltage of -90 mV to voltages between -100 and +40mV in 10 mV steps. **B** Average ± SEM of the IV relationships of inward currents (circles) and steady-state outward current (triangles) from WT (black, n = 14) and *Tmem16b* KO (blue n = 13) OSNs. Values were taken at the negative peak and at the end of voltage step as indicated by the symbols in A (p = 0.73 unpaired t-test for inward current; p = 0.64 U-test for outward current). Scatter dot plot with averages ± SEM of input membrane resistance (**C**) and zero-current membrane potential (**D**) of WT and *Tmem16b* KO OSNs (for R_m_ n = 12-13 p= 0.24, U-test; for zero-current potential n=9-10 p= 0.28, U-test). Representative whole-cell recording of OSNs from WT (**E**) and *Tmem16b* KO mice (**F**) stimulated with IBMX (1 mM) at the indicated holding potential. **G** Scatter dot plot with averages ± SEM of ratios of the IBMX-induced current at -90 mV and at -50 mV (n = 10 for WT and n = 9 for KO; p = 2.16*10^-5^ U-test).

Although we found no differences in the biophysical properties we measured, we previously showed that in *Tmem16b* KO OSNs the transduction current is dramatically reduced (Pietra et al., 2016). Here, we confirmed that when stimulated for 1 s with the phosphodiesterase inhibitor IBMX (1 mM). We set the holding potential to -50 mV, the equilibrium potential for Clˉ with our solutions (see Methods) and the transduction currents in WT and KO OSNs were similar in amplitude (Fig. 1E-F) between WT and *Tmem16b* KO. Switching the holding potential to -90 mV greatly amplified the transduction current revealing the contribution of the Clˉ currents (Fig. 1E tangerine trace) in the WT. In the KO, there were no differences between the transduction currents recorded at different holding potentials (Fig. 1F) and the ratio between current amplitudes at -90 mV and -50 mV is close to 1 (Fig. 1G) in *Tmem16b* KO while it was increased 3-fold in the WT. Altogether, these results indicate that the absence of Clˉ current in OSNs do not change either their biophysical properties or the basic voltage-gated currents. On the other hand, the transduction current is greatly reduced in the OSNs from *Tmem16b* KO mice confirming the significant contribution by the Clˉ current.

### The Clˉ channel shapes kinetic parameters of the odorant response

To address how TMEM16B contributes to the odorant-induced response and its kinetics, we stimulated WT and *Tmem16b* KO OSNs with an odor mixture (100 µM of cineole and acetophenone each) and recorded the responses with the suction pipette technique. WT OSNs displayed the typical rapid rise of the suction current upon odorant exposure and decayed during the 1 s stimulation to decline gradually thereafter back to baseline (Fig. 2A). OSNs lacking TMEM16B responded with a similar delay, reached their peak current faster, but generated a much smaller maximal current as expected. The response current also declined during the 1 s stimulation and had a faster decline back to baseline at the end of stimulation compared to WT. To display and address action potential firing, we filtered odorant responses at a wider bandwidth (0 – 5 kHz). WT OSNs generated only a few action potentials at the onset of stimulation (Fig. 2B) while KO OSNs could generate longer action potential trains (Fig. 2C).

**Figure 2.**
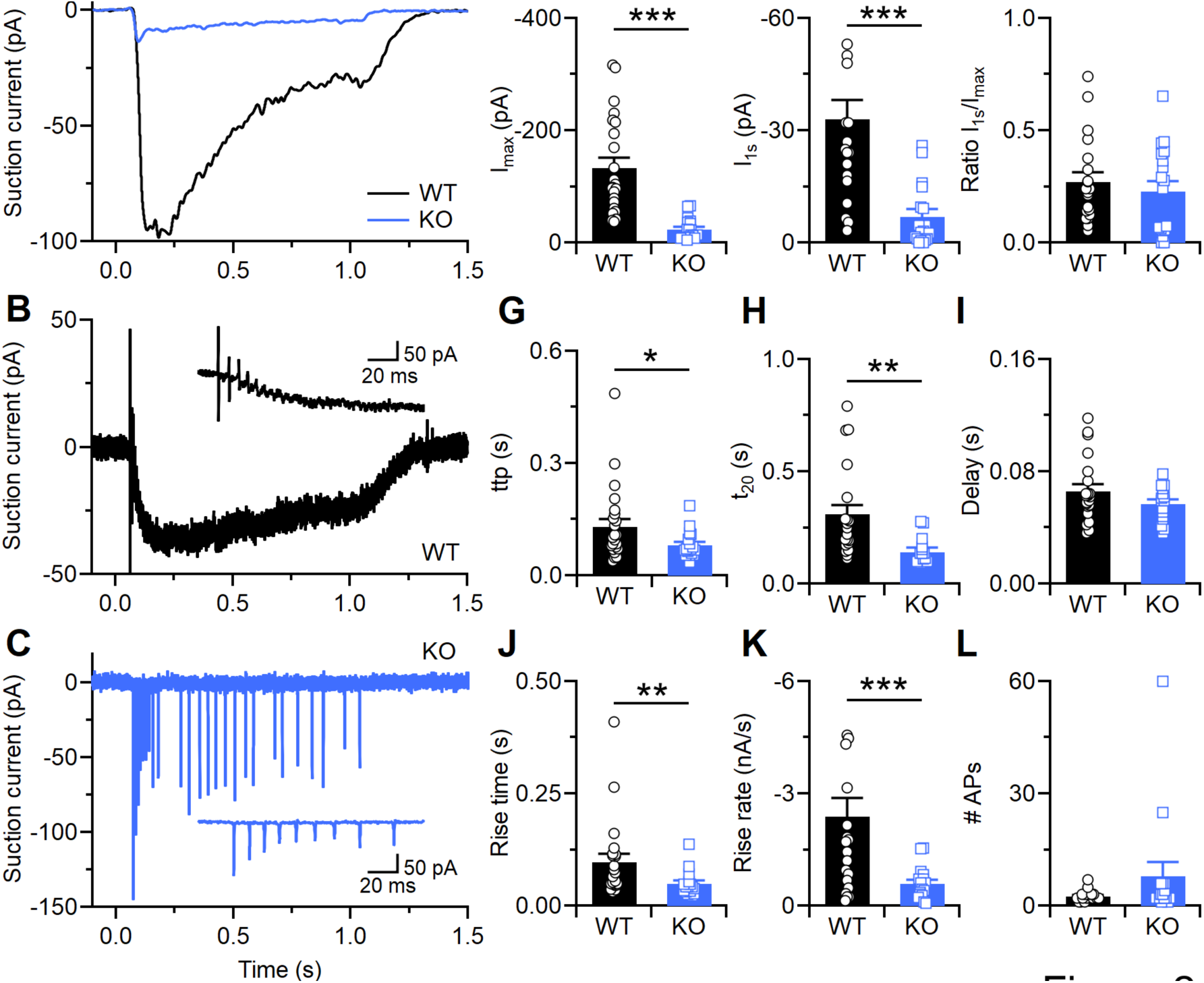
The lack of TMEM16B affects odorant response kinetics. **A** OSNs were stimulated with 100 µM of cineole and acetophenone each for 1 s as indicated by the solution monitor on top. Current responses were recorded using the suction pipette technique and filtered 0 – 50 Hz to display only the receptor current. **B** & **C** display WT and KO OSN responses filtered at the wide bandwidth of 0 – 5 kHz to also display action potentials. **D** Peak current I_max_ elicited by the odorant response **E** current I_1s_ at the end of the odorant stimulation **F** Ratio of I_1s_ and I_max_. **G** time to peak (ttp), the time to reach Imax following odor onset **H** t_20_, the time for the current to fall to 20 % of I_1s_ from the end of stimulation **I** Response delay measured as the time taken for the first action potential to be generated following odor onset **J** Rise time, ttp minus delay, **K** Rise rate of the receptor current, Imax divided by rise time, **L** number of action potentials generated during the 1 s stimulation. n = 20 – 23 OSNs for WT and 14 - 18 for KO. * p < 0.05, ** p < 0.01, *** p < 0.001, t test. Bars indicate mean and error bars are SEM.

On average, KO OSNs only generated a maximal current of around 20 % of the WT OSNs (Fig. 2D). The current at 1 s was similarly reduced (Fig. 2E). This is reflected in a similar ratio of the current at 1 s when normalized to its maximal current (Ratio I_1s_/I_max_, Fig. 2F). As the KO lacks the Clˉ current, this suggests that this decline in current response during the 1 s stimulation mostly depends on a reduction of the current through the CNG channels, reduced Ca^2+^ influx and hence reduced activation of the Clˉ channel. Thus, the Clˉ channel itself might contribute little to this form of adaptation. Lack of Clˉ current curtailed the time the current required to reach its peak response (time to peak, ttp), from 130 ± 21 ms in WT to 81 ± 8 ms the KO (Fig. 2G), indicating that while the current through the CNG channel might reach its maximum quickly, continuing, although decreasing, influx of Ca^2+^ can continue to increase ciliary Ca^2+^ and further increase the Clˉ current thereafter. At the termination of the response, the time required for the current to fall to 20 % of the current at 1 s is also shortened from 310 ± 40 ms in the WT to 140 ±20 ms in the KO (Fig. 2H), confirming the role of the Clˉ current in determining the response duration. Interestingly, the response delay, measured as the time between the onset of stimulation and the generation of the first action potential was unchanged in the KO compared to the WT and therefore seems to be mostly determined by the initial speed of activation of the CNG channel (Fig. 2I). The rise time (Fig. 2J), the duration from the response delay to the time to peak, as well as the rise rate (Fig. 2K), the peak current divided by the rise time, was larger in the WT compared to the KO. The number of action potentials generated in response to odor stimulation was not statistically different in WT and KO (Fig. 2L), although the KO in several cells showed an extreme prolongation of the action potential train.

### The absence of Clˉ channels greatly reduce response amplitude without altering odorant sensitivity

We studied the role of TMEM16B in the odorant-induced dose response by exposing OSNs to an odorant mix of 3, 10, 30 and 100 µM of cineole and acetophenone each. In WT OSNs, the increase in the stimulus concentration led to increasingly larger receptor current responses (Fig. 3A), which also became longer. In KO OSNs, responses also increased, although at a much-reduced current level and the responses terminated quickly at the end of stimulation (Fig. 3B). Fig. 3C compares the average responses across WT and KO OSNs, with again the KO OSNs showing much smaller responses. We also asked if the sensitivity of KO OSNs might be reduced. We normalized the dose response relation of each OSN to its response at 100 µM and averaged across all OSNs for WT and KO respectively (Fig. 3D). The two dose response relations showed no relative shift to each other, suggesting that odorant sensitivity was largely unaltered in KO OSNs. The numbers of action potentials fired across the odorant concentrations used showed that KO OSNs generated more action potentials compared to the WT (2-way ANOVA, Fig. 3E).

**Figure 3.**
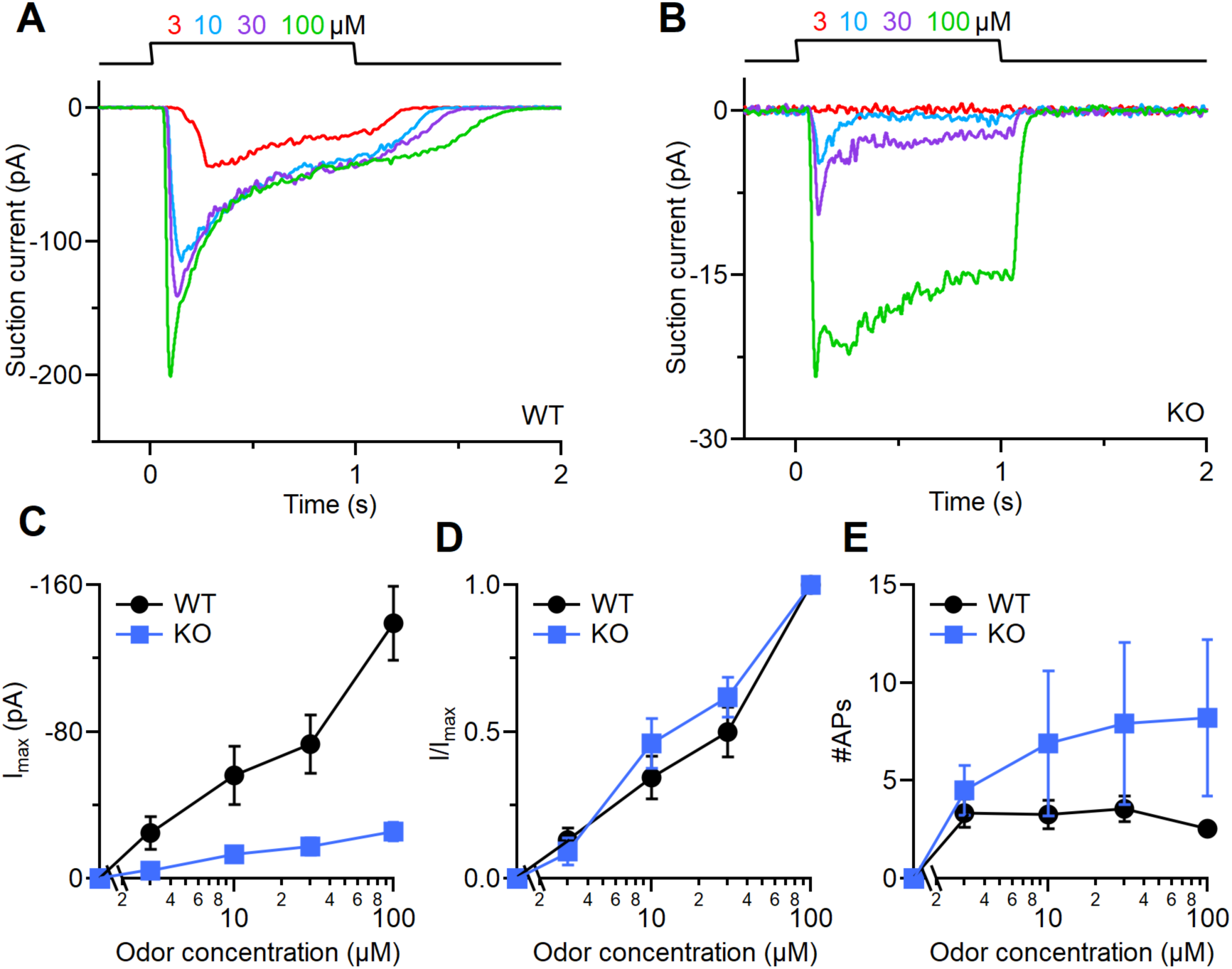
The lack of TMEM16B does not alter the OSN odor sensitivity. **A** WT and **B** *Tmem16b* KO OSNs were stimulated with increasing odorant concentrations at the indicated concentration (mixture of cineole and acetophenone) for 1 s. Responses were recorded using the suction pipette technique and filtered 0 – 50 Hz. **C** Dose response relation of the maximal peak current I_max_. **D** The dose response of each OSN was normalized to its response at 100 µM. n = 17 - 20 for WT and 13 – 16 for KO. Data points are mean ± SEM. **E** Number of action potentials generated in WT and KO OSNs, which was statistically higher in the KO (2-way ANOVA, F(1,107) = 23.96, p = 0.0106.

### Clˉ current contribution to the recovery from adaptation

As the odorant responses terminated more quickly in the KO OSNs, and the speed of response termination is a major determinant in the speed of recovery from odorant-induced adaptation (Stephan et al., 2011; Dibattista and Reisert, 2016), we performed double pulse experiments where OSNs were exposed twice for 1 s with increasing interpulse interval in between the two stimulations. Fig. 4A shows such an experiment with an interval duration of 0.5 s for a WT OSN. In response to the first stimulation, the OSN generates a large response with action potentials also being generated at the onset of the response. At the end of the first stimulation, the current declined slowly and had not yet reached zero levels upon the beginning of the second stimulation. While the OSN still generated an, although small, increase in receptor current, it failed to generate action potentials at this point, most likely as voltage-gated Ca^2+^ and Na^+^ channels were still inactivated from the preceding depolarization (Trotier, 1994; Pietra et al., 2016). In contrast (Fig. 4B), while the responses of the KO OSN were much smaller, the response to the first stimulation terminated quickly, allowing the OSN to reach its basal level of current and hence hyperpolarization to generate action potentials again when stimulated for the second time. To analyze the responses, we plotted the ratio of the second current response divided by the first against the interpulse interval (Fig. 4C). In both the WT and the KO, the response to the second stimulation progressively decreased with decreasing interpulse interval, although the relative response size in the KO remained larger across all interpulse intervals, indicating that recovery from adaptation occurs on a faster timescale in the KO compared to the WT. To understand how action potential firing is affected, we calculated the chance of action potentials being generated when exposed to the second odor pulse. For long interpulse intervals, both WT and KO OSNs generated action potentials very reliably, but at interpulse intervals of 1 s and shorter, WT OSNs became less reliable in generating action potentials (Fig. 4D). For the KO, a drop in reliable action potential generation was only observed at the shortest interpulse interval of 0.25 s (Fig. 4D).

**Figure 4.**
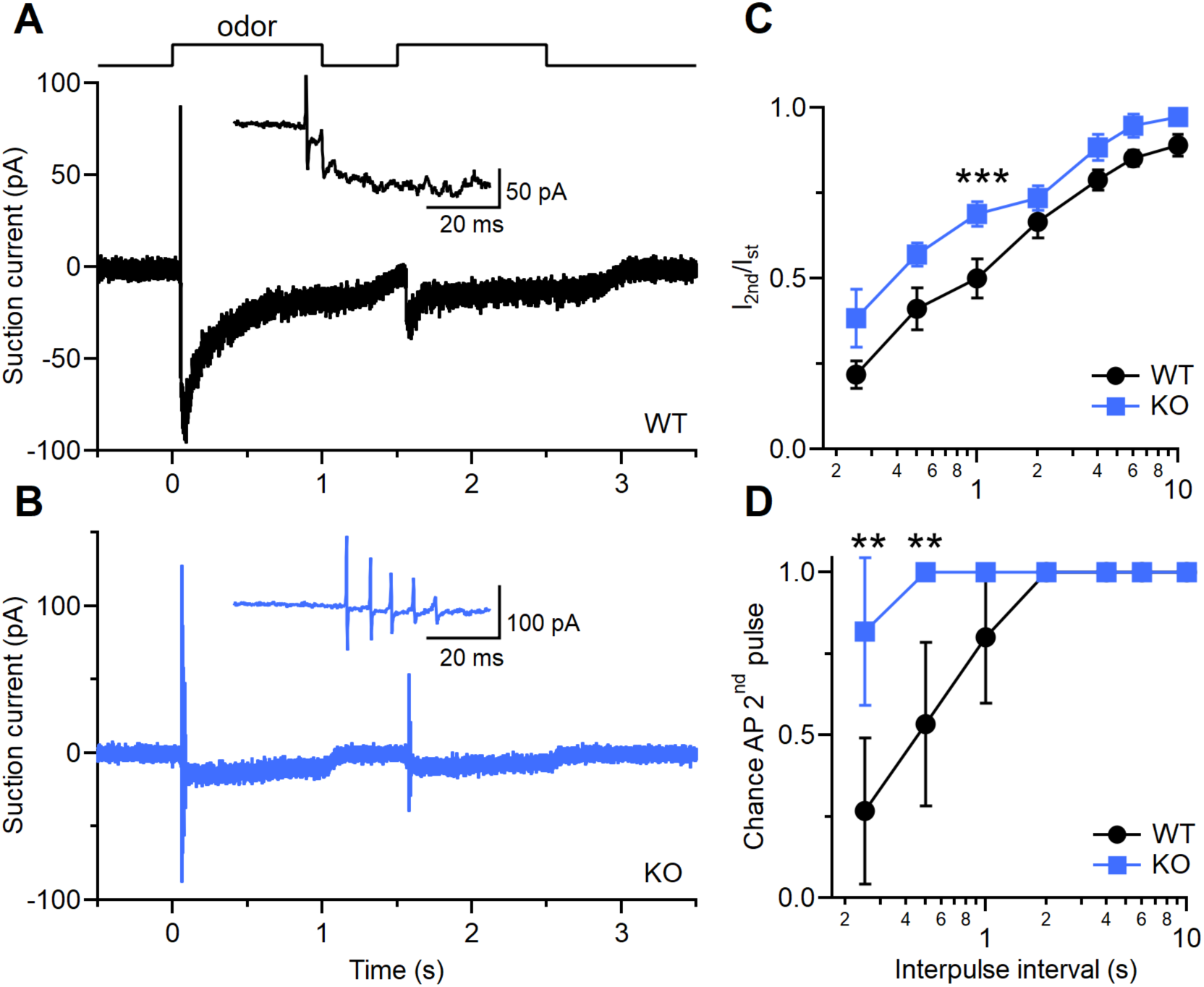
The lack of TMEM16B fastens the recovery from adaptation. Double pulse experiments to determine response recovery following adaptation. **A** A WT and **B** *Tmem16b* KO OSN were stimulated twice for 1 s with an interpulse time of 0.5 s. Odorants were cineole and acetophenone at 100 µM each, currents were recorded with the suction pipette technique and filtered at 0 – 5 kHz. Note that both WT and KO OSN generated action potentials to the first stimulus, but the WT OSN failed to generate action potentials in response to the second stimulus in contrast to the KO OSN. **C** For increasing interpulse times, the peak current of the second response was normalized to the first response. Data are mean ± SEM. WT and KO are significantly different, 2-way ANOVA, F(1,160) = 23.96, p = 2.3 10^-6^. **D** Chance that OSNs fire an action potential in response to the second odorant exposure, data points are mean ± 95% confidence interval, ** p < 0.01, Fisher exact test. n = 11 – 15 OSNs for WT and KO.

### Odorant responses to different frequencies of stimulation

Given the previous result, we asked how WT and KO OSNs respond to a change in stimulation frequency from low (2 Hz, resembling breathing at rest) to a higher (5 Hz, resembling a more sniff-like) frequency. A WT OSN (Fig. 5A) reasonably tracked a 2 Hz stimulation pattern and generated action potentials in response to most stimulations, but, in this case failed to respond to the second and eighth stimulation. Its response reliability dropped even further when stimulated at 5 Hz thereafter. The KO OSN (Fig. 5B) generated action potentials to every stimulation at 2 Hz, but did so with less reliability at 5 Hz. Fig. 5C compares the percent of stimulations with fired action potentials at 2 Hz between WT and KO across odorant concentrations. The KO could maintain a high, around, and above 75 % firing reliability across the odorant concentrations, while in the case of the WT, with the exception of the lowest concentration, it stayed at around 50 %, which was statistically lower compared to the KO. At 5 Hz, both WT and KO showed low, around 25 %, firing reliability. Thus, while at lower stimulation frequencies KO OSNs are more reliable, this difference is not observed any longer at 5 Hz.

**Figure 5.**
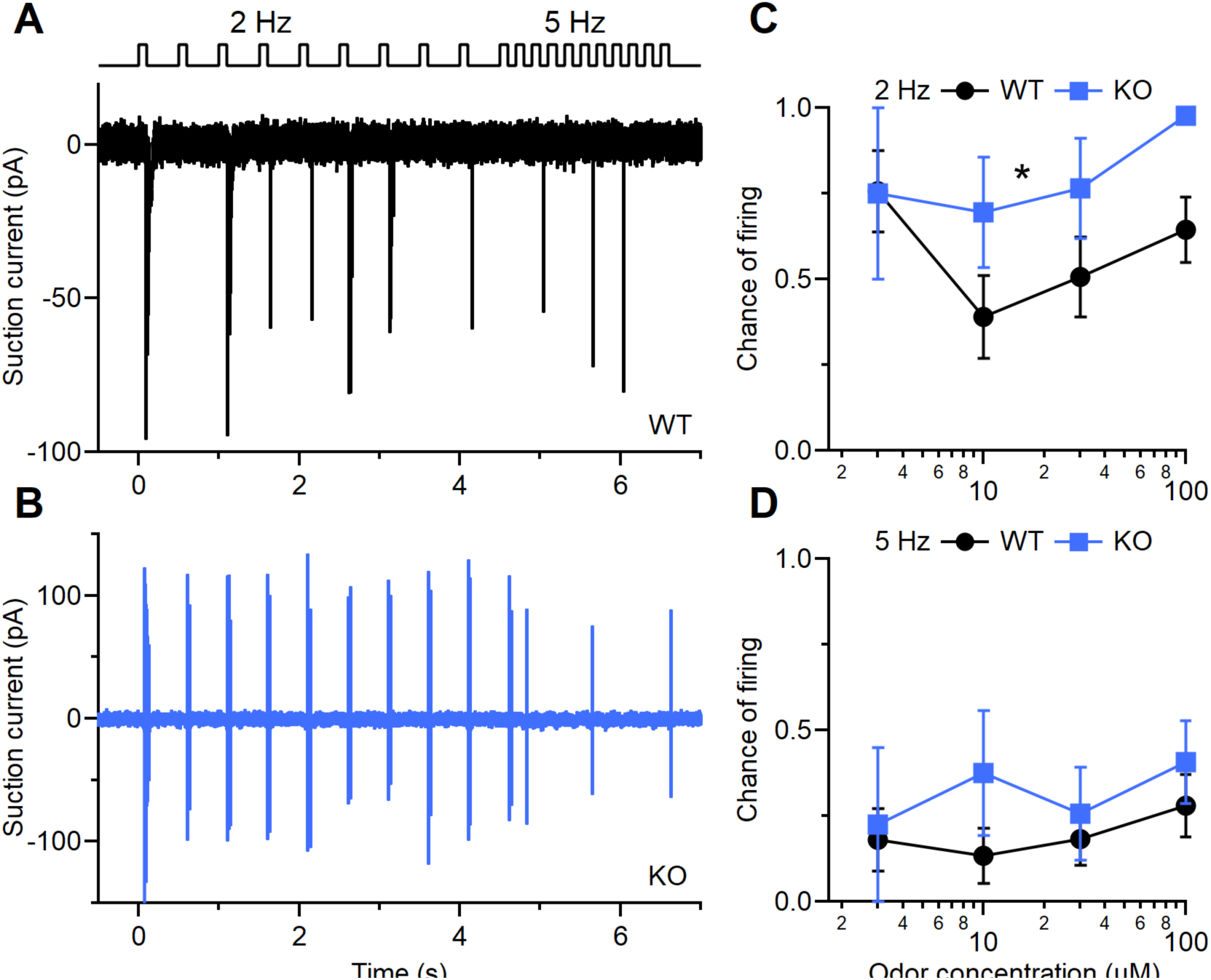
Odorant responses to 2 and 5 Hz stimulations. **A** A WT and **B** *Tmem16b* KO OSN were stimulated at 2 Hz followed by 5 Hz as indicated by the solution monitor at the top (100 ms stimulus duration, 100 µM cineole and acetophenone each). Currents were recorded with the suction pipette technique at 0 – 5 kHz. **C** and **D** Reliability of WT and *Tmem16b* KO OSNs to generate action potentials at each 2 Hz and at 5 Hz stimulation. At 2 Hz, WT and KO are statistically different, 2-way ANOVA, F(1,80) = 67.3, p = 0.02. n = 5 - 20 for WT and 4 – 14 for KO. Data are mean ± SEM.

We sought to extend the previous sets of data by performing experiments using slices from olfactory epithelia and recorded using the loose-patch configuration. Similar to the suction pipette recordings, this configuration does not alter the intracellular milieu and additionally avoids any isolation-induced artifact. We applied five repetitive 1 s odor stimulation with an interpulse interval of 0.5 s, a similar recovery interval as used during the 2 Hz stimulation in the suction pipette recordings (see Fig. 5). The WT OSNs fired action potentials almost exclusively at the first 1 s odor application while the *Tmem16b* KO were firing at each stimulation (Fig. 6A). The number of action potentials and duration of the action potential train was altered in the KO, being respectively higher and longer than in the WT (Fig. 6C; number of APs WT 3.7 ± 0.8 n = 7, KO 13 ± 3 n = 7, p = 0.0047 U-test; duration WT 144 ± 79 ms n = 7, KO 670 ± 176 ms n = 7, p = 0.017 U-test). During the first stimulation, the WT decreased its firing frequency and fell silent quickly, while the KO progressively reduced its firing frequency but continued to fire throughout the 1 s stimulation. Interestingly, the lack of silencing did not influence the ability of fire action potentials at each stimulation in the KO. We quantified an adaptation index as a ratio between the number of action potentials in response to the n^th^ stimulation and the number of action potentials at the first. The WT OSNs nearly completely stopped firing as the action potentials in the stimulations following the first were very few or even absent (Fig. 6 E). The KO had a number of action potentials that was about 60 % of the first stimulus and it stabilized at around 50 % across subsequent stimulations (Fig. 6E). In summary, using this stimulation paradigm we conclude that OSNs from *Tmem16b* KO follow the stimulation with higher fidelity (see also Fig. 5). Because of our perfusion and experimental slice set-up, we could not shorten the interpulse interval further. When we increased the interval to 1 s (Fig. 6B), we could again observe an increase in the number and action potential train duration in the KO (Fig. 6D), but the adaptation index was similar between WT and KO (Fig. 6F).

**Figure 6.**
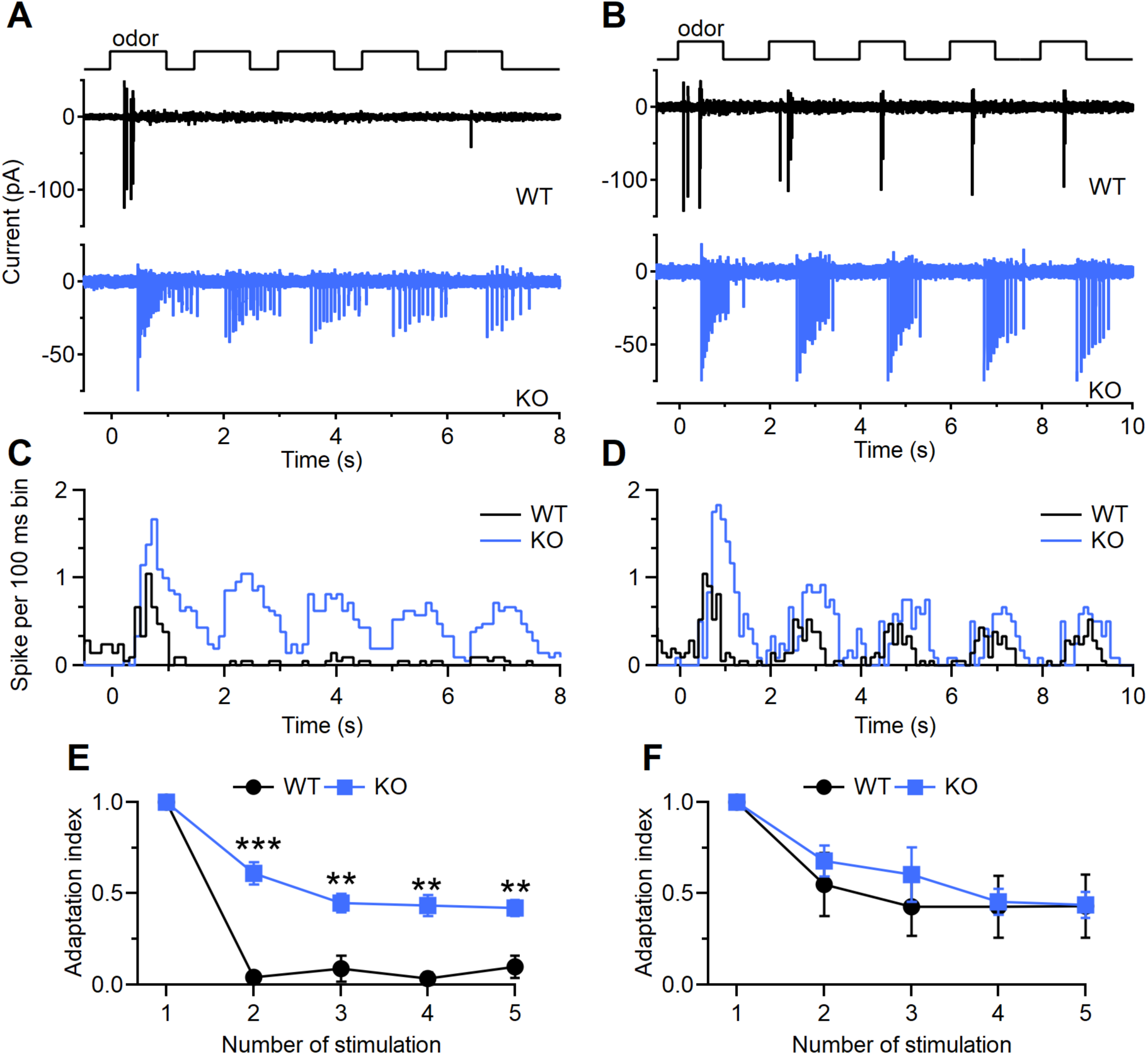
The lack of TMEM16B fastens the recovery from adaptation after odor stimulation in loose-patch recording. Representative loose-patch recordings from OSNs from WT and *Tmem16b* KO mice stimulated with an odor mix for 5 times with an interpulse interval of 0.5 s (**A**) or 1 s (**B**). (**C** and **D**) Peristimulus histograms showing the sum of number of spikes for all the cells normalized by the number of cells (bin = 100 ms) for experiments in **A** and **B** (n = 7 for WT and KO). (**E** and **F**) Adaptation index (ratio of action potentials evoked by the n^th^ stimulation and the number of action potentials evoked by first stimulation) from experiments in **A** and **B** (n = 7 for WT and n = 4 KO; *** p < 0.001 **p < 0.01 U-test).

We also performed the same type of stimulation protocol but instead of an odorant mix we used IBMX. The responses in term of number of action potential and action potential train duration were qualitatively similar to those obtained with the odorant mix (Fig. 7A-F; number of action potentials wt 6.4 ± 1.6 n = 12, KO 14.9 ± 2.6 n=12, p = 0.0075 U-test; duration WT 142 ± 47 ms n=12, KO 819 ± 103 ms n = 12, p = 9*10^-5^ U-test). With an interpulse interval of 0.5 s, in the WT few action potentials were reemerging from adaptation in the later pulses thus increasing the adaptation index compared to that observed with the odorant mix (compare Fig. 7A with 6A). The differences between the adaptation indexes of the WT and the KO were also present especially at the 2^nd^ and 3^rd^ IBMX applications (Fig. 7E). All differences between WT and KO were no longer observed when we increased the stimulus interval to 1 s (Fig. 7B-F).

**Figure 7.**
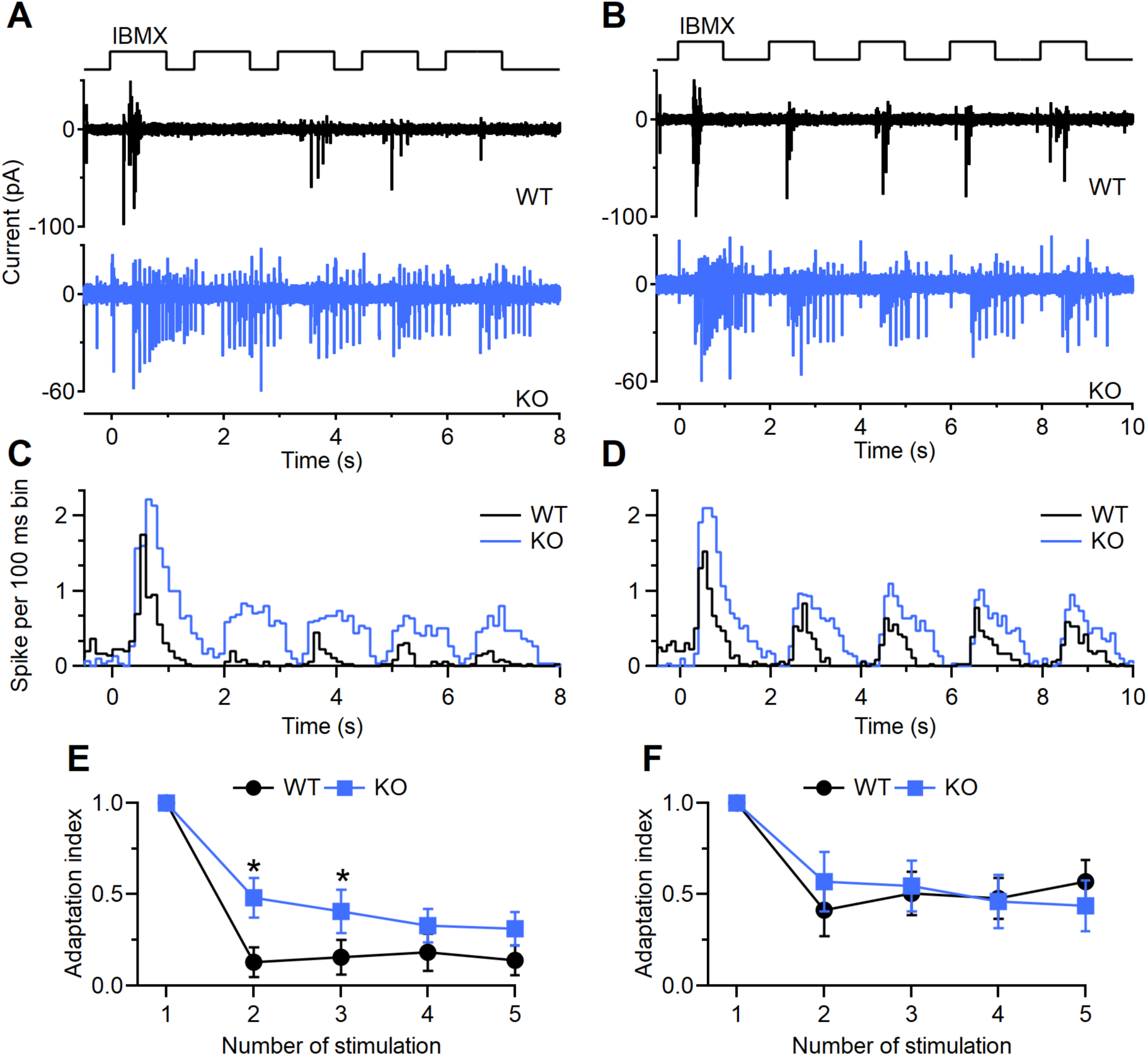
The lack of TMEM16B fastens the recovery from adaptation after IBMX stimulation in loose-patch recording. Representative loose-patch recordings of OSNs from WT and *Tmem16b* KO mice stimulated with 1 mM IBMX 5 times with an interpulse interval of 0.5 s (**A**) or 1 s (**B**). (**C** and **D**) Peristimulus histograms showing the sum of number of spikes for all the cells normalized by the number of cells (bin = 100 ms) for experiments in **A** and **B** (n = 12 for WT and n = 10 for KO). (**E** and **F**) Adaption index (ration of APs evoked by the n^th^ stimulation and the number of APs evoked by first stimulation) from experiments in **A** and **B** (n = 12 for WT and n = 10 for KO; *p<0.05 U-test).

In summary, OSNs from *Tmem16b* KO have altered adaptation properties failing to modulate the number of action potentials and spike train duration according to the stimulation frequency.

## Discussion

The events that lead to the generation of the receptor potential in OSN cilia are triggered by the binding of an odorant to the OR and shaped by the currents generated via the cationic CNG channel first and anionicTMEM16B current thereafter. It has long been established that the Cl^-^ current provides high gain, low noise amplification of the receptor current (Lowe and Gold, 1993; Kleene, 2008; Li et al., 2018), while the CNG channel’s contribution to shaping the time course and overall sensitivity, in particular during repetitive stimulation, has received less attention (Pifferi et al., 2012; Dibattista et al., 2017, 2024).

The Cl^-^ opens following the opening of the CNG channel and Ca^2+^ influx, which is the main source of ciliary Ca^2+^ rises (Nakamura and Gold, 1987; Firestein et al., 1991; Lowe and Gold, 1993; Schild and Restrepo, 1998; Kleene, 2008; Boccaccio et al., 2021). Hence its activation is closely coupled to the activation kinetics of the CNG channel. This is consistent with the observed similar response delay in both WT and *Tmem16b* KO OSNs and suggests that the response delay is mainly dependent on the speed of generation of the second messenger cAMP and the ensuing activation of the CNG channel. The decline of the odorant response during a 1 s stimulation was similar in WT and KO OSNs, suggesting that this form of adaptation is mostly driven by the gating of the CNG channel and thus, either via desensitization of the CNG channel or a reduction in available cAMP, either by reduced cAMP synthesis or increased ciliary cAMP hydrolysis by PDE (Boccaccio et al., 2006, 2021; Cygnar and Zhao, 2009; Dibattista and Reisert, 2016).

Response termination following cessation of odorant stimulation requires the closure of the Cl^-^ channel. Which itself depends on the closure of the CNG channel and cessation of Ca^2+^ influx and additionally on the removal of ciliary Ca^2+^, the latter uncoupling the Cl^-^ channel from CNG channel kinetics. It has long been known that when inhibiting Ca^2+^ extrusion by removing extracellular Na^+^, thus abolishing the driving force of Na^+^-dependent Ca^2+^ extrusion, the response is greatly prolonged and carried by a sustained Ca^2+^-activated Cl^-^ current (Jung et al., 1994; Reisert and Matthews, 1998, 2001; Antolin and Matthews, 2007). The main molecular determinant of Ca^2+^ extrusion is the K^+^-dependent Na^+^/ Ca^2+^ exchanger 4 (NCKX4) and *Nckx4* knockout OSNs display prolonged response termination by up to several seconds (Stephan et al., 2011). Here, in the *Tmem16b* KO, we observed a faster response termination than in the WT, consistent with the view that it is the Cl^-^ current that determines the duration of the response. Interestingly and importantly, the rate of termination also determines recovery from adaptation during repetitive odorant stimulation (Stephan et al., 2011; Dibattista and Reisert, 2016). It is a key cellular behavior for OSNs to fine-tune their dynamic response across various input levels, both in stimulation magnitude as well as stimulation frequency, the latter varying with the sniffing frequency of the mouse (Firestein et al., 1990; Ghatpande and Reisert, 2011). In particular, adaptation to repeated stimuli manifests as a reduction in the current amplitude of the response to the second odorant stimulus with respect to the first (Boccaccio et al., 2006; Song et al., 2008; Stephan et al., 2011). This difference in amplitude of the responses is reduced when the time interval between stimuli is increased; complete recovery of the response is seen for a sufficiently long interval. In the *Tmem16B* KO the response recovery is faster than in the WT, meaning that for a given interpulse interval, the response to the second stimulation in a *Tmem16b* KO regains a larger amplitude normalized to the first when compared to the WT. This could be explained by at least two not-mutually exclusive ways. The Cl^-^ channel itself might undergo inactivation during prolonged increased Ca^2+^ levels, for which there is evidence that this is the case (Reisert et al., 2003; Ponissery Saidu et al., 2013). Alternatively, as overall current levels and therefore cellular depolarization is changed in the *Tmem16b* KO, this will also alter Ca^2+^ influx and ciliary Ca^2+^ levels that could lead to altered adaptation of the transduction cascade. As stated above, response termination is important to determine adaptation and *Nckx4*4 KO OSNs showed that NCKX4 is required to lower intraciliary Ca^2+^ during and after odorant stimulation, allowing the transduction cascade to recover from adaptation (Stephan et al., 2011).

While the Cl^-^ current amplifies the odorant response, it remains unclear if OSNs alter their odorant sensitivity in the absence of the Cl^-^ channel. We performed odorant dose-response experiments and, as expected, WT odorant responses were larger compared to the KO. But normalization of the responses to their maximal current revealed that WT and KO dose-response relations overlap and are not shifted relative to each other. This suggests that the sensitivity of OSNs is mostly determined by events upstream of the Cl^-^ channel and that odorant sensitivity does not seem to be affected by the lack of the Cl^-^ current. These results seem to disagree with the idea that the Cl^-^ current may non-linearly amplify OSN responses as the Ca^2+^-activated Cl^-^ current is highly cooperative (Lowe and Gold, 1993; Li et al., 2016, 2018). A potential explanation of the observed differences is that previous work was performed in voltage-clamped OSNs, while in our experiments, the voltage was free to vary, which might limit the amount of current an OSN can generate (Lowe and Gold, 1993). Experiments using suction pipette recordings were performed at room temperature which makes it difficult to compare with ours (Li et al., 2018). A potential caveat here is that OSNs we recorded from were randomly selected and we used a mix of cineole and acetophenone as the stimulating odorants. Thus, we have taken an “odor-centric” approach to the question, but have not addressed this question in an “OR-centric” perspective, where one would record from OSNs expressing known ORs.

The APs constitute the output of OSNs that are being sent to the olfactory bulb. OSNs increase their firing rate upon increasing odorant concentration. The number of APs, though, first increases monotonically with increasing concentration, but reaches a plateau or even decreases at higher odorant concentrations (Reisert and Matthews, 2001; Rospars et al., 2008). Interestingly, OSNs fire APs only in the very early phase of the odorant response in isolated OSNs, generating usually between 2 or 3 APs, rarely 5 or more due to a rapidly declining AP amplitude brought on by inactivation of voltage-gated Na^+^ and Ca^2+^ channels (Trotier, 1994; Pietra et al., 2016). Only when recording from OSNs of *Tmem16B* KO this number increase to 8 (or more) during dose-response experiments. Triggering APs also depends on the input resistance of the OSNs, which is overall high and small currents are sufficient to elicit AP firing. We did not detect any substantial changes either in voltage-gated currents or in input resistance between WT and KO. Therefore, similar input resistance of OSNs from WT and KO guarantee that the current needed to trigger APs would substantially be the same. The increase in the number of APs that we observed in the *Tmem16B* KO may be due to the slower rise rate (slope) of the receptor current in the KO rather than its overall time course.

The termination of the receptor current is important for odorant perception during repeated stimulation. In our double pulse paradigm, for short interpulse intervals of 0.5 s, around 50% of WT OSNs failed to generate APs in response to the second stimulation but reliably improve when the interpulse interval is lengthened. An increase in stimulation frequency (shorter interpulse interval) progressively abolished AP generation. *Tmem16b* KO OSNs were able to fire APs reliably (close to 100%) even at the shortest (0.25 s) interpulse interval. We investigated OSN firing behavior in more detail by shortening both stimulus duration and the interpulse interval so that our protocol mimicked normal breathing (2 Hz) and higher-frequency sniffing (5 Hz). We observed that *Tmem16b* KO OSNs generated APs more faithfully than WT in response to 2 Hz stimuli. Thus, similar to the results in the double pulse experiments, lack of Cl^-^ current apparently allows OSNs to track stimulations more reliably. At 5 Hz, both WT and KO OSNs tracked stimulations only poorly, suggesting that the faster response termination seen in the KO can now not compensate any longer for the higher stimulation frequency. Nevertheless, it might seem counterintuitive why the lack of the Cl^-^ channel yields an OSN that generates more APs more reliable during repeated stimulation, an observation consistent with the observed larger responses when imaging from OSN axon terminals in the OB of *Tmen16b* KO mice (Zak et al., 2018). Or in other words, why the addition of a Cl^-^ current in OSNs seems to abolish encoding of odor stimuli during repeated stimulation? An interesting notion here is the concept of adaptive filtering (Ghatpande and Reisert, 2011). As mice are able to modulate their breathing frequency and hence the stimulation frequency of their own OSNs, they can also alter the olfactory information they receive with an increase in breathing frequency suppressing the sensation of present odorants, but still enable the perception of new odorants. Our data suggests that the Cl^-^ could be an important, if not the determining, component in such a mechanism. They might also explain an apparent discrepancy between behavioral deficits reported for *TMEM16B* KO mice. In a Go/Nogo experiments, mice show no behavioral deficits (Billig et al., 2011), while in food search tasks, KO mice perform worse compared to WT (Pietra et al., 2016; Neureither et al., 2017). A potential explanation is that in the Go/Nogo task only the first inhalation might suffice to perform the odor identification task, which can be conducted in as short as 200 ms (Uchida and Mainen, 2003; Abraham et al., 2004; Rinberg et al., 2006), while the search task required repeated odor sampling over time, which is heavily impacted as we show here.

In summary, most of the response characteristics of the odorant response in OSNs depend on the Cl^-^ current. Therefore, it not only amplifies the odorant response, but also shapes response kinetics acting synergistically with NCKX4 in determining the optimal rate of termination. By doing so, they control cellular properties that are directly linked to response termination such as adaptation to repeated stimulation.

## METHODS

Mice were handled in accordance with the guidelines of the Italian Animal Welfare Act and European Union guidelines on animal research, under a protocol approved by the ethic committee of SISSA or in accordance with methods approved by the Animal Care and Use Committees of the Monell Chemical Senses Center (conforming to National Institutes of Health guidelines).

For suction pipette experiments, heterozygous *Tmem16b* breeder mice were obtained from T. Jentsch, Max-Delbrück-Centrum für Molekulare Medizin, Berlin, Germany (Billig et al., 2011) and bred to obtain WT and *Tmem16b* KO mice. Adult mice of either sex were euthanized by CO_2_ inhalation and decapitated. The head was split sagitally along the midline and the olfactory epithelium was removed. The tissue was stored in glucose-containing Ringer solution at 4°C until further use. Ringer contained in mM: 140 NaCl, 5 KCl, 1 MgCl_2_, 2 CaCl_2_, 0.01 EDTA, 10 HEPES, and 10 glucose. The pH was adjusted to 7.5 with NaOH. Odorants, cineole and acetophenone (Sigma), were dissolved into Ringer at the stated concentrations.

Odorant-induced responses from isolated OSNs were recorded using the suction pipette technique as previously described (Lowe and Gold, 1991; Ponissery Saidu et al., 2012; Dibattista and Reisert, 2018). OSNs were isolated by placing olfactory epithelium in a Ringer-filled Eppendorf, followed by gentle vortexing. OSNs were allowed to settle in the recording chamber and subsequently sucked into the tip of the recording pipette. Currents were recorded with a Warner PC-501A patch clamp amplifier, digitized using a Power1401 II A/D converter and Signal acquisition software (Cambridge Electronic Design). Currents was acquired twice, once filtered at 0 -50 Hz to display the receptor current and once 0 – 5 kHz to resolve the fast biphasic action currents that drive the action potentials. The sampling frequency was 10 kHz.

OSNs were exposed to odorants using the Perfusion Fast-Step solution changer (Warner Instrument Corporation). The tip of the recording pipette with the sucked OSN was transferred quickly across the interface of two neighboring solution streams, which were heated to mammalian body temperature of 37°C (Matthews, 1999).

### Preparation of acute slices of mouse olfactory epithelium

Acute coronal slices of olfactory epithelia of P0 - P4 mice were obtained with a method similar to previously described by Henriques et al. (2019) and Agostinelli et al. (2021). Mice at this age are suitable for use since TMEM16B is already express at P0 - P4 (Maurya and Menini, 2014; Maurya et al., 2015). For these experiments, we used *Tmem16b* KO mice kindly provided by Prof. Lily Jan (University of California, San Francisco, USA, (Zhang et al., 2017)). After the mouse was sacrificed, the head without the skin was dissected and embedded in 3 % low grade agar prepared in Ringer’s solution once the solution had cooled to 38°C. After solidification the agar block was fixes in a glass Petri dish filled by oxygenated Ringer’s solution and a vibratome was used to cut coronal slices of 300 μm thickness (Vibratome 1000 Plus Sectioning System). The slices were kept in cold oxygenated Ringer’s solution until use.

### Electrophysiological recordings from OSNs and stimuli presentation

Slices were viewed with an upright microscope (BX51WI, Olympus) equipped with a 40X water immersion objective with an additional 2X auxiliary lens. The recording chamber was continuously perfused with oxygenated Ringer’s solution at room temperature. Extracellular solutions were exchanged through an eight-in-one multibarrel perfusion pencil connected to a ValveLink 8.2 pinch valve perfusion system (Automate Scientific). The bath was connected with an Ag/AgCl reference electrode through a 3 M KCl agar bridge. OSNs were identified by their morphology. The experiments were performed at room temperature (20 - 25 °C). Patch pipettes were pulled from borosilicate capillaries (WPI) with a Narishige PC-10 puller.

In the whole-cell voltage clamp configuration experiments the pipette had a resistance of 4 - 7 MΩ when filled with the intracellular solution. We used an intracellular solution composed of (in mM) 80 KGluconate, 60 KCl, 2 MgATP, 1 EGTA, and 10 HEPES (adjusted to pH 7.2 with KOH) to measure the voltage-gated currents and the passive properties of the OSNs. The voltage-gated current was measured in voltage clamp mode, holding the potential at -90 mV and applying a voltage step protocol from -100 mV to + 40mV with 10 mV increment. In voltage clamp mode, we also evaluated the membrane input resistance. We held the potential of the cell at -90 mV and gave a negative pulse of 20 mV. In these conditions, the voltage-gated currents are not activated (Dibattista et al., 2008). In current clamp mode (without injecting current), we measured the resting membrane potential. To isolate the Clˉ current contribution, we used an intracellular solution composed of (in mM) 13 KCl, 132 KGluconate, 4 MgCl_2_, 0.5 EGTA, and 10 HEPES (adjusted to pH 7.2 with KOH). In this way, the reversal potential for Clˉ, after on-line correction for the liquid junction potential, corresponds to the holding potential of -50 mV. In this condition the current is mainly mediated by CNG channels.

Extracellular recordings from the soma of OSNs were obtained in the loose patch configuration using a pipette with a resistance of 3 - 4 MΩ when filled with Ringer’s solution. During the experiments the seal resistances were 30 - 50 MΩ. Loose patch experiments were made in voltage clamp mode with a holding potential of 0 mV. The recordings were performed using a MultiClamp 700B amplifier controlled by Clampex 10.6 via Digidata 1550B (Axon Instrument, USA). Data were low pass filtered at 2 kHz and sampled at 10 kHz.

The cells were stimulated with a mix of odorants composed of heptanal, isoamyl acetate, acetophenone, cineole and eugenol. Each odorant was prepared as a 5 M stock in dimethyl sulfoxide (DMSO) and on the day of experiments was diluted in Ringer’s solution to reach the final concentration of 100 µM each. We used 3-isobutyl-1-methylxanthine (IBMX). IBMX is a phosphodiesterase (PDE) inhibitor which induces an increase of the cAMP concentration in a subpopulation of OSNs. It was prepared weekly and used at the concentration of 1 mM dissolved directly into Ringer’s solution without the use of DMSO. At this concentration IBMX, elicits responses with kinetics similar to those induced by odorants (Reisert et al., 2007).

## Data analysis

IgorPro software (Wavemetrics) was used for data analysis and to make the figures. Data are given as mean ± SEM. Normal distribution of the data was tested with Shapiro-Wink test. The homogeneity of the variance was tested using Levene’s test. If the data are normally distributed and with the same variance statistical significance was determined using unpaired homoscedastic Student’s t test or 2-way ANOVA. If data presented with a different variance, an unpaired different variance Student’s t test was used. For not normally distributed data, the Mann-Whitney U-test was used. P values of p < 0.05 were considered statistically significant. The loose patch recordings were filtered offline with a high pass filter at 5 Hz to correct for baseline drifts. Spikes were detected using an event detection algorithm setting an arbitrary threshold and then were confirmed by shape examination.

## Acknowledgements

*TMEMKO* mice were kindly provided by Drs. G. Billig and T. Jentsch and Dr. L. Jan. The research was supported by NIH grant R01DC016647 (awarded to J.R.). Monell’s animal facility renovations were supported by NIH grant G20-OD020296 for infrastructure improvement.

